# A eukaryote without tRNA introns

**DOI:** 10.1101/2025.06.17.660180

**Authors:** Ambro van Hoof, Tokiko Furuta, Swathi Arur

**Affiliations:** Department of Microbiology and Molecular Genetics. The University of Texas Health Science Center at Houston Houston TX 77030; Department of Genetics. The University of Texas MD Anderson Cancer Center Houston, TX 77030

## Abstract

One of the striking characteristics of eukaryotic genomes is the presence of three types of introns: spliceosomal introns, tRNA introns, and a unique intron in the *XBP1* mRNA. Exceptional eukaryotic genomes that lack spliceosomal or *XBP1* introns have been described. However, tRNA introns and the tRNA endonuclease that is required for their splicing are thought to be universal in eukaryotes. The introns in three tRNAs are widely conserved across Metazoa: Tyr-GUA, Ile-UAU and Leu-CAA. This study shows that some nematode species have lost the introns in Tyr-GUA and Ile-UAU tRNAs, and one species, *Levipalatum texanum*, completely lacks tRNA introns. The loss of the intron from Leu-CAA tRNA is accompanied by an unusual A-C mismatched basepair in the anticodon stem loop and a triplication of a tRNA deaminase that could potentially restore basepairing. These changes may be an adaptation to the loss of the intron. *L. texanum* also lacks the tRNA endonuclease, one of two enzymes required for tRNA splicing. The other key enzyme in tRNA splicing, tRNA ligase, is bifunctional and is also required for *XBP1* mRNA splicing. *L. texanum* retains tRNA ligase and the *XBP1* intron. This eukaryote without tRNA introns has the potential to be a valuable tool for disentangling the functions of tRNA splicing, *XBP1* splicing and tRNA modification enzymes, and is the only animal known to have lost one of the three intron types.

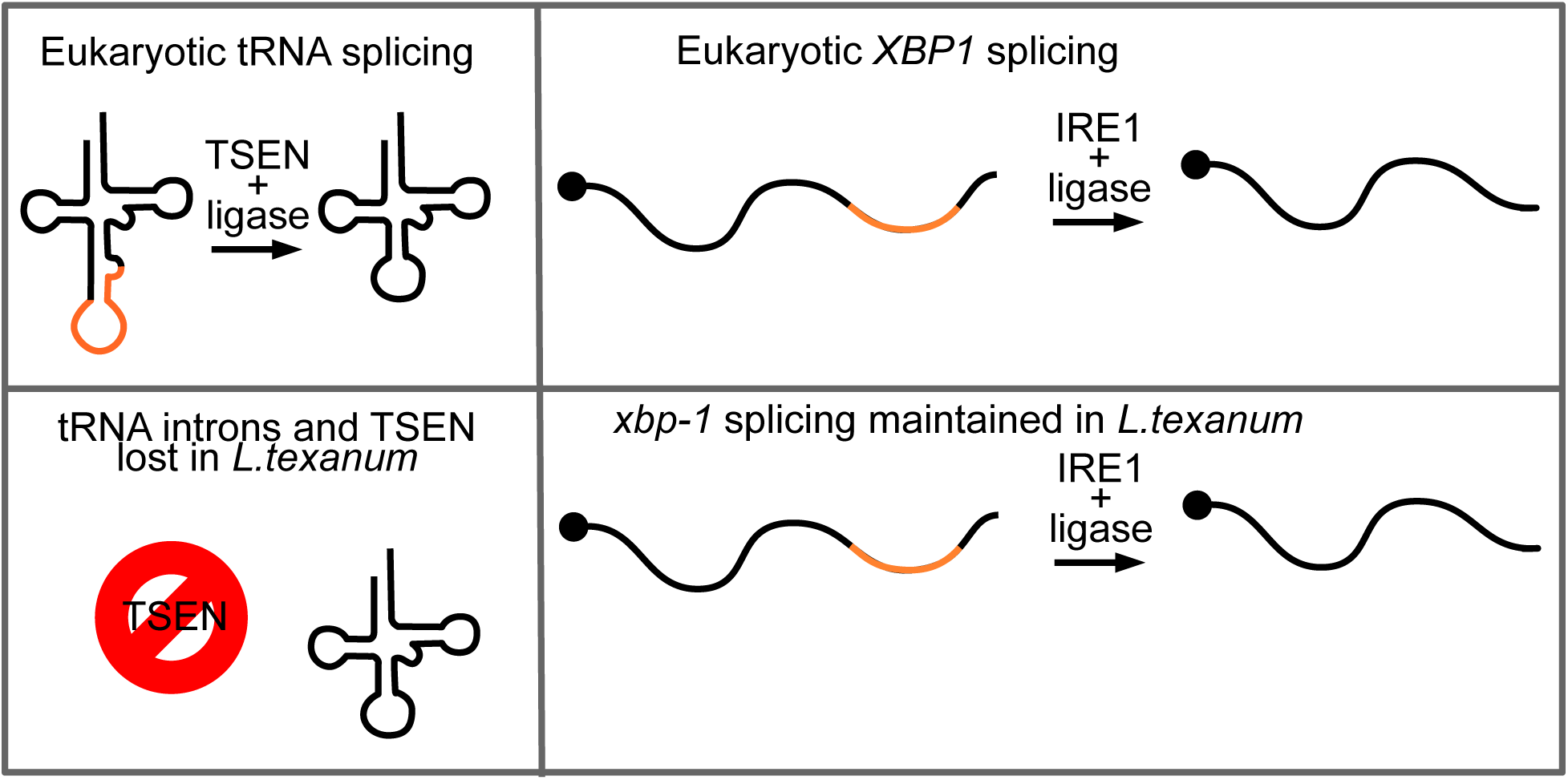

**SIGNIFICANCE:** tRNA introns are thought to be a universal feature of eukaryotic genomes, including all animal genomes. We identified a species of roundworm, *Levipalatum texanum*, that completely lacks tRNA introns. *L. texanum* also lacks the tRNA splicing endonuclease, one of two enzymes required for the removal of introns from tRNAs. The second enzyme involved in this process, tRNA ligase, has a second function: the splicing of a single mRNA. Interestingly, this mRNA splicing reaction is conserved. *L. texanum* has an usual leucine tRNA and a three copies of a tRNA modifying enzyme that may compensate for the absence of the intron. These findings highlight that although complex RNA processing is a hallmark of eukaryotes, individual reactions may be absent in specific eukaryotes.

## INTRODUCTION

Most eukaryotes have three types of introns in their nuclear genomes, and the three intron types appear to be completely unrelated to each other. The most well-known are the spliceosomal introns in mRNAs and other RNA polymerase II transcripts. These introns are removed by a complex molecular machine, the spliceosome, whose snRNA subunits are essential for both intron recognition and catalysis. Metazoan genomes typically contain many thousands of spliceosomal introns, although other eukaryotes can have only a few. A second type of intron is found in the mRNA encoding the transcription factor XBP1/Hac1 and its orthologs. This intron is removed by an endoribonuclease, IRE1, followed by ligation of the two exons by an RNA ligase (1, 2). This *XBP1* splicing is initiated during the unfolded protein response (UPR), which activates IRE1. The UPR splicing mechanism appears dedicated to the unique *XBP1/HAC1* intron. Finally, some tRNAs contain introns that are removed by tRNA splicing endonuclease (TSEN) followed by ligation of the two exons (3, 4). The *XBP1* and tRNA splicing pathways are initiated by different RNases but use the same RNA ligase (5).

All three types of introns are widespread in animals, fungi, and plants. Spliceosomal and tRNA introns are also present in even earlier diverging eukaryotes, such as trypanosomes and the malaria parasite, suggesting that these two intron-types were present in the ancestor of all eukaryotes. On the other hand, the IRE1-dependent splicing of *XBP1* appears absent from early diverging eukaryotes (6), suggesting that it may be the most recent addition to the spectrum of eukaryotic introns. Although the three types of introns are a conserved feature of eukaryotic genomes, their function is not fully clear, especially for tRNA introns. It was recently suggested that after tRNA introns are released from the pre-tRNA, they can regulate mRNAs by base pairing to them (7). A more established role of tRNA introns is that some of them are required for proper tRNA modification (8, 9). Several tRNA modification enzymes appear to act on the intron-containing precursor and are unable to modify the spliced tRNA (8, 9).

Although the three splicing pathways are generally well conserved, there are exceptions. For example, the model fungus *Schizosaccharomyces pombe* appears to lack an IRE1-dependent intron and an *XBP1* ortholog (10). Interestingly, *S. pombe* has maintained IRE1, but this endonuclease only appears to function in mRNA degradation (11). Likewise, the spliceosomal introns and the spliceosome appear to be lost from some microsporidia, which are fungi with minimized genomes (12–15). In contrast, tRNA introns and the tRNA splicing machinery are thought to be a universal feature of eukaryotic genomes (5), The number of tRNA introns in a eukaryotic genome can vary from one to hundreds but a genome with no tRNA introns has not been reported (https://gtrnadb.ucsc.edu/index.html) (16).

To the best of our knowledge all previously analyzed animal genomes contain all three intron types. We report that the genome of the nematode *Levipalatum texanum* lacks tRNA introns. Furthermore, it lacks the tRNA splicing endonuclease that is central to tRNA splicing. *L. texanum* is related to the more well-characterized *Pristionchus pacificus* and *Caenorhabditis elegan*s species of nematode. *P. pacificus* is used as a model organism to study evolutionary developmental biology, and *C. elegans* is one of the premier model organisms in biology. Unlike its close relatives *P. pacificus* and *C. elegans*, *L. texanum* was described very recently and is thus a largely unstudied species (17). It was first isolated in 2006 in Texas, and the existing genome assembly and transcriptome are from an independent 2011 isolate from Virginia (18). Its biology remains uncharacterized, and its genome has been used mainly to better understand *P. pacificus* biology.

During analysis of tRNA intron distributions across eukaryotes, we noted that a draft genome of *L. texanum* lacked any tRNA introns or the tRNA endonuclease required to splice them. The draft genome consisted of 33,427 contigs. To exclude the possibility that we missed an intron, we generated a new 408 contig genome assembly from long PacBio HiFi reads, eliminating >98% of the gaps and increasing the continuity of the genome by 200-fold (N50 of 2,003 kbp instead of 8.6 kbp). Comparison of the genome of *L. texanum* with related nematodes revealed that an early ancestor nematode of the order Rhabditida contained introns in three tRNAs (Tyr tRNA with a GUA anticodon, Ile tRNA with a UAU anticodon and Leu tRNA with a CAA anticodon; hereafter tY(GUA), tI(UAU) and tL(CAA), respectively. We observe that within this order, the introns from tY(GUA) and tI(UAU) were lost in a subset of the species within the Diplogasteridae family. Within this family, the tL(CAA) intron was also lost in the *L. texanum* lineage after it diverged from all other nematodes with sequenced genomes. Our finding that the *L. texanum* genome does not contain the endonuclease needed for tRNA splicing suggests that this enzyme became nonessential after loss of the introns. Importantly, these data suggest that the three classes of introns are not universal across eukaryotes. In yeast, TSEN has a second essential function that is unrelated to tRNA splicing (19, 20). The observation that TSEN has been lost from *L. texanum* suggests that this second TSEN function is not conserved across eukaryotes. Finally, tRNA ligase and the IRE1/tRNA ligase-dependent splicing of *XBP1* are conserved in *L. texanum*. These observations suggest that *L. texanum* is uniquely suited to study tRNA biology in the absence of tRNA introns as well as the role of the tRNA ligase complex in *XBP1* splicing in the absence of confounding effects of tRNA splicing.

## RESULTS AND DISCUSSION

### The tyrosine tRNA genes of the nematode *Pristionchus pacificus* and other Diplogasteroidea lack introns

The genomic tRNA database (https://gtrnadb.ucsc.edu/index.html) is an authoritative database of tRNA sequences across organisms (21). The tRNAs are initially predicted by tRNAscan-SE 2.0 (16), and for some genomes (including model organisms) further manually annotated to eliminate low confidence predictions and pseudogenes. In an effort to understand the distribution of tRNA introns, we analyzed the 575 eukaryotic genomes in version 21 of GtRNAdb (9). This analysis showed that all but one of those genomes had an intron in tY(GUA). The one exception in GtRNAdb was the nematode *P. pacificus*. The *P. pacificus* genome contains 1021 tRNA genes predicted with high confidence that encode 48 different tRNAs (Supplemental Informa9on). The tY(GUA) tRNA is encoded by 29 genes in *P. pacificus*, and all 29 lack an intron (Figure 1). However, the eight *P. pacificus* tL(CAA) genes are each interrupted by an intron. In addition, the *P. pacificus* genome carries possible introns in individual predicted genes for tG(GCC) (1 intron-containing and 24 intronless genes), tV(CAC) (1 intron-containing and 15 intronless genes), and tL(CAG) (1 intron-containing and 16 intronless genes). Thus, while *P. pacificus* is an exceptional eukaryote without introns in tY(GUA), this species is still predicted to require tRNA splicing to produce a full set of tRNAs needed to translate all 61 sense codons: In the absence of splicing, UUG codons cannot be translated, which account for 6% of all of its leucine codons (https://www.kazusa.or.jp/codon/cgi-bin/showcodon.cgi?species=54126).

**Figure 1:**
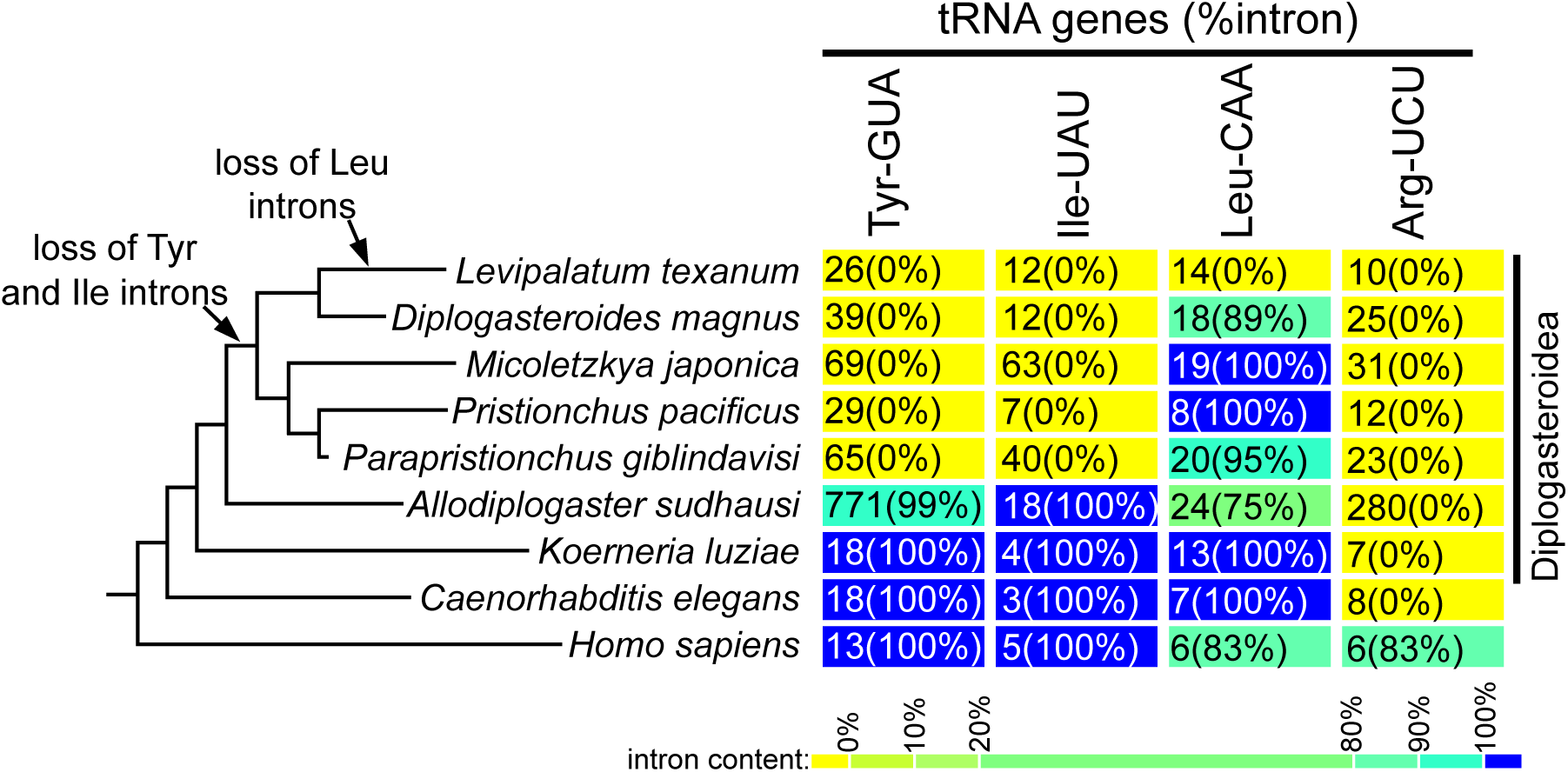
tRNA intron loss in Diplogasteroidea nematodes. Shown is a phylogenetic tree of *L. texanum*, related nematodes, and human on the left. The tRNA gene and intron count for the four indicated tRNAs is shown on the right. For each species and each tRNA, the number of tRNA genes is indicated. The percentage of them that contain an intron is indicated in parenthesis and by the background color/heatmap. Yellow indicates no introns. Blue indicates all tRNA genes contain an intron. *C. elegans* and humans have a typical set of metazoan tRNA introns. In the Diplogasteroidea, five species have lost the introns in tY(GUA) and tI(UAU), and *L. texanum* has additionally lost the introns in tL(CAA) and has no tRNA introns. Branch lengths in the tree are arbitrary.

Although GtRNAdb is a valuable source of tRNA gene sequences, not all sequenced eukaryotic genomes are included, and the completeness of the included genome sequences is not easily verified. To explore the idea that one or more intron-containing tY(GUA) genes might have been missed in *P. pacificus*, the genome sequences of closely related nematodes that are not included in GtRNAdb were retrieved from Genbank and analyzed by tRNAscan-SE 2.0 (16) (Figure 1). *P. pacificus* is a member of the same order as the much better studied *C. elegans*, but within this order is in a different superfamily, the Diplogasteroidea. Most of the Diplogasteroidea genomes also lacked introns in their tY(GUA) genes. Specifically, our tRNAscan-SE 2.0 analysis indicated that the genomes for *Parapristionchus giblindavisi*, *Micoletzkya japonica*, *Diplogasteroides magnus* and *L. texanum* contained between 26 and 69 predicted tY(GUA) genes, and importantly, all genes are intronless (Figure 1). In contrast, in the earlier diverging (22) *Allodiplogaster sudhausi* genome, 99% of the tY(GUA) genes contained an intron. Therefore, tY(GUA) appears to have lost its intron in a subset of Diplogasteroidea after their divergence from *A. sudhausi*, making this clade of nematodes the first group of eukaryotes described without an intron in tY(GUA).

The intron in tY(GUA) genes in animals, fungi, and plants is required to modify the GUA anticodon to GΨA (23–27). In addition, in yeast this intron is required to convert A58 to ^m1^A58 (9). However, these modifications are not essential for yeast or human viability (9, 28–30). Thus, either these modifications do not occur in Diplogasteroidea, or they are generated in an intron-independent manner.

### The genome of the nematode *Levipalatum texanum* lacks tRNA introns

To determine whether tY(GUA) was the only tRNA missing an intron in the Diplogasteroidea or whether this phenomenon was more broadly spread in this superfamily, we analyzed the other tRNAs and compared them to tRNAs from human and *C. elegans*. Upon tRNAscan-SE analysis of Diplogasteroidea genomes, we observed something striking: the *L. texanum* genome lacked introns in all of its tRNA genes. However, the genome sequences analyzed were draft genomes assembled from short (Illumina) whole genome sequencing reads. For example, for the initial analysis of the *L. texanum* genome, we used an assembly (https://www.ncbi.nlm.nih.gov/datasets/genome/GCA_014805385.1/) consisting of 33,427 contigs with an N50 of 8.6 kbp. Thus, there was a possibility that the genome assembly was missing some tRNA genes. To eliminate this possibility, we needed a higher quality assembly of the *L. texanum* genome with fewer gaps. Thus, we generated long PacBio HiFi reads and assembled them with hifiasm (31) into a more complete and contiguous genome of 205,346,893 bp consisting of 408 contigs with an N50 of 2,003,394 (meaning that half of the sequence is present in contigs larger than 2 Mbp). This newly assembled genome is available from Genbank under accession number PRJNA1267722 and is used for the analysis described below. Importantly, this improved assembly confirmed that tRNA genes in *L. texanum* are indeed devoid of introns (Figure 1), similar to the conclusions drawn from the lower quality Illumina based GCA_014805385.1 assembly.

According to the GtRNAdb, introns are prevalent in metazoan tY(GUA), tI(UAU), tL(CAA), and tR(UCU) genes but essentially absent from the other metazoan tRNAs (9, 21). Consistent with this metazoan pattern, both human and *C. elegans* have introns in their tY(GUA), tI(UAU), and tL(CAA) genes, and human additionally has introns in tR(UCU) genes. (Figure 1; https://gtrnadb.ucsc.edu/genomes/eukaryota/Hsapi38/; https://gtrnadb.ucsc.edu/genomes/eukaryota/Celeg11/). Our tRNAscan-SE analysis of the earliest diverging Diplogasteroidea, *K. luziae* and *A. sudhausi*, showed that they have very similar tRNA intron distributions to *C. elegans*. However, the other five Diplogasteroidea each lack introns in the tI(UAU) and tY(GUA) tRNA genes (Figure 1). Thus, while tY(GUA) and tI(UAU) genes have introns in almost all Metazoa, including *C. elegans*, these introns were lost in the ancestor of these five related species. Three of the Diplogasteroidea retained the intron in all of their tL(CAA) genes.

*D. magnus* lost the tL(CAA) intron in two of its 18 tL(CAA) genes. Strikingly, *L. texanum* lost the intron in all 14 of its tL(CAA) genes. tRNAscan-SE 2.0 predicted no high confidence tRNA genes with introns for *L. texanum*. These analyses of Diplogasteroidea genomes show that a common ancestor of *P. giblindavisi*, *P. pacificus, M. japonica*, *D. magnus* and *L. texanum* lost tI(UAU) and tY(GUA) introns (Figure 1). We conclude that the tL(CAA) intron was the last one lost from *L. texanum*, after its divergence from *D. magnus* and all other nematodes.

### The *L. texanum* genome lacks the genes for tRNA splicing endonuclease

If tRNA introns were indeed lost from the *L. texanum* genome, was the tRNA splicing machinery also lost? Eukaryotic tRNA introns are removed from pre-tRNA by a heterotetrameric TSEN (3, 32–34). TSEN consists of two catalytic subunits, named TSEN2 and TSEN34 in human, and two noncatalytic subunits, TSEN54 and TSEN15 in human (33, 34). Similar subunits are conserved in fungi (Sen2, Sen34, Sen54, and Sen15 in the yeast *S. cerevisiae*). The two catalytic subunits have higher sequence similarity across eukaryotes and are readily identified across Metazoa. The noncatalytic subunits are more diverged in sequence and not as readily identified. Furthermore, TSEN15 is the smallest subunit (<20KDa), which makes it more difficult to identify orthologs in other eukaryotes. For example, *C. elegans* has annotated *tsen-2*, *tsen-34* and *tsen-54* genes that are the orthologs of *TSEN2*, *TSEN34* and *TSEN54*, respectively, but does not have an annotated *tsen-15*, and we could not identify a candidate *C. elegans tsen-15* by homology searches (https://wormbase.org/resources/gene_class/tsen-01--10). Although, TSEN has not been characterized in *C. elegans*, TSEN-2, TSEN-34 and TSEN-54 appear to be similar to the orthologous TSEN subunits from other eukaryotes and are likely functionally equivalent.

We used the human and *C. elegans* TSEN subunit sequences to search both genome assemblies of *L. texanum*. As a control, the closest relative with tRNA introns, *D. magnus*, was analyzed similarly. TBLASTN searches of the *D. magnus* genome and BLASTP searches of its annotated proteins readily identified obvious candidate genes for TSEN-2, TSEN-34 and TSEN-54, but despite exhaustive effort, *L. texanum* TSEN orthologs could not be identified (Table 1; Supplemental Figure S2), consistent with the absence of tRNA introns in this species. We also used the protein sequences predicted from the *D. magnus* TSEN genes as queries of the *L. texanum* genome and predicted proteins. Although these species are closely related, this search did not detect any *L. texanum* TSEN orthologs.

**Table 1:** tRNA endonuclease subunit genes are absent from the *Levipalatum texanum* genome.

|  | human | *C. elegans* | *D. magnus* | UTH_Ltex_0 genome assembly |  |  |  |
| --- | --- | --- | --- | --- | --- | --- | --- |
|  |  |  |  | gene | contig | start | end |
| tRNA endonuclease | *TSEN2* | *tsen-2* | 08070g021655 | none found |  |  |  |
|  | *TSEN34* | *tsen-34* | 00304g003500 | none found |  |  |  |
|  | *TSEN54* | *tsen-54* | 00100g001689 | none found |  |  |  |
| TSEN-associated | *CLP1* | *clpf-1* | 07871g021449 | *clpf-1* | 174 | 107,521 | 109,758 |
| tRNA deaminases | *ADAT1* | none found | none found | none found |  |  |  |
|  | *ADAT2* | *JC8.4* | 08507g022124 | *adat-2* | 29 | 965,410 | 967,041 |
|  | *ADAT3* | *Y47D3A.14* | 00091g001593 | *adat-3a* | 42 | 836,125 | 832,741 |
|  |  |  |  | *adat-3b* | 42 | 911,297 | 906,829 |
|  |  |  |  | *adat-3c* | 42 | 868,275 | 860,339 |
| intron-dependent tRNA modification enzymes | *NSUN2* | *nsun-2* | 00735g006105 | *nsun-2* | 138 | 208,228 | 199,297 |
|  | *PUS1* | *pus-1* | 00130g002035 | *pus-1* | 211 | 60,266 | 63,011 |
|  | *PUS7* | *B0024.11* | 00036g000729 | *pus-7* | 288 | 25,661 | 28,308 |
|  | *TRMT61* | *W02A11.1* | 00191g002600/<br>00191g002601** | *trmt-61* | 32 | 1,127,345 | 1,122,272 |
|  | *TRMT6* | *trmt-6* | 03866g015564 | *trmt-6* | 203 | 54,786 | 52,797 |
| tRNA ligase complex | *RTCB* | *rtcb-1* | 00729g006080 | *rtcb-1* | 108 | 146,782 | 148,888 |
|  | *DDX1* | *Y55F3BR.1* | 00216g002818 | *ddx-1* | 126 | 34,187 | 28,969 |
|  | *FAM98B* | *tag-322* | 03174g014111 | *tag-322* | 63 | 166,980 | 164,322 |
|  | *CGI-99* | *E02H1.5* | 03780g015391 | *cgi-99* | 167 | 47,824 | 49,034 |
| ligase activation | archease | *C43H8.1* | 00354g003848 | *arch-1* | 143 | 128,199 | 127,146 |
| UPR | *XBP1* | *xbp-1* | 00164g002358 | *xbp-1* | 53 | 1,068,412 | 1,069,497 |
|  | *IRE1* | *ire-1* | 06002g019109/<br>02462g012347/<br>02462g012345* | *ire-1* | 5 | 2,736,759 | 2,715,233 |

\*the *ire-1* genes in the Illumina based genome assemblies of *L. texanum* and *D. magnus* are split between two scaffolds with a sequence gap in one of the introns. In both cases, the intact gene could be assembled based on RNAseq reads that span this intron. Three separate parts of the *D. magnus ire-1* gene are annotated as individual genes. The reduced number of gaps in the HiFi Pacbio based genome assembly resulted in the *L. texanum ire-1* gene being contained in a single contig.
\*\* The *D. magnus trmt-61* gene is annotated as two separate genes, but RNAseq clearly shows that the two parts are spliced together and form one functional gene.

The lllumina *L. texanum* genome assembly consists of 33,427 contigs, and thus it is possible that some genes are missed in this assembly. This provided the main motivation to assemble our higher quality genome sequence. However, as discussed below, many genes of interest could readily be identified in the initial 33,427 contigs. We were able to identify *pus-1, trmt-61*, and *ire-1* orthologs in the first draft genome, even though these genes have an intron that spans two scaffolds. The more contiguous HiFi assembly indeed confirmed that the two *pus-1, trmt-61*, and *ire1* fragments were adjacent in the genome, but we did not detect any tRNA splicing related genes in the PacBio HiFi assembly that are missing from the Illumina assembly. In addition, the *D. magnus* genome sequence contains 42,037 contigs and thus is far worse than our assembly with 408 contigs. However, despite this lower quality, we readily detected the *D. magnus* genes for TSEN subunits (Table 1). It thus seems highly unlikely that all of the genes for tRNA splicing endonuclease are in the gaps in the HiFi *L. texanum* genome assembly. Therefore, we conclude that the *tsen-2*, *tsen-34* and *tsen-54* genes were lost from the *L. texanum* genome after its divergence from *D. magnus*.

In addition to the TSEN subunits common to eukaryotes, Metazoa have a fifth TSEN subunit named CLP1, or CLPF-1 in *C. elegans* (33). The function of CLP1 in tRNA splicing is unclear, but CLP1 has additional functions, including in the cleavage and polyadenylation machinery that generates mRNA 3’ ends. The *L. texanum* genome encodes a clear ortholog of CLPF-1 (Table 1). Since TSEN subunits are absent, CLPF-1 is likely maintained for its mRNA processing functions. Overall, these analyses show that *L. texanum* lost both tRNA introns as well as the tRNA splicing endonuclease complex required to remove introns from pre-tRNAs.

### The intronless tRNA Leu CAA genes of *L. texanum* encode an unusual A-C basepair

To explore the possibility that the *L. texanum* tL(CAA) evolved to somehow compensate for the absence of the intron, the sequences for each of the tL(CAA) tRNAs from the *L. texanum* genome were used to generate a multiple sequence alignment (depicted as a sequence logo in Figure S3). The same analysis was performed for the tL(CAA) tRNAs of *C. elegans*, *P. pacificus* and *D. magnus*, all of which contain introns (Figure S3). The spliced tL(CAA) tRNAs from *C. elegans*, *P. pacificus*, and *D. magnus* were highly similar, including an identical anticodon stem loop (Figure 2A) despite these nematodes diverging 186 million years ago (https://timetree.org/). This anticodon stem loop is also identical to the human tL(CAA). In contrast to this conservation, the intron loss of the *L. texanum* tL(CAA) gene was accompanied by many other changes, including in the anticodon loop, the anticodon stem, and the acceptor stem (Figure 2A and Figure S3).

**Figure 2:**
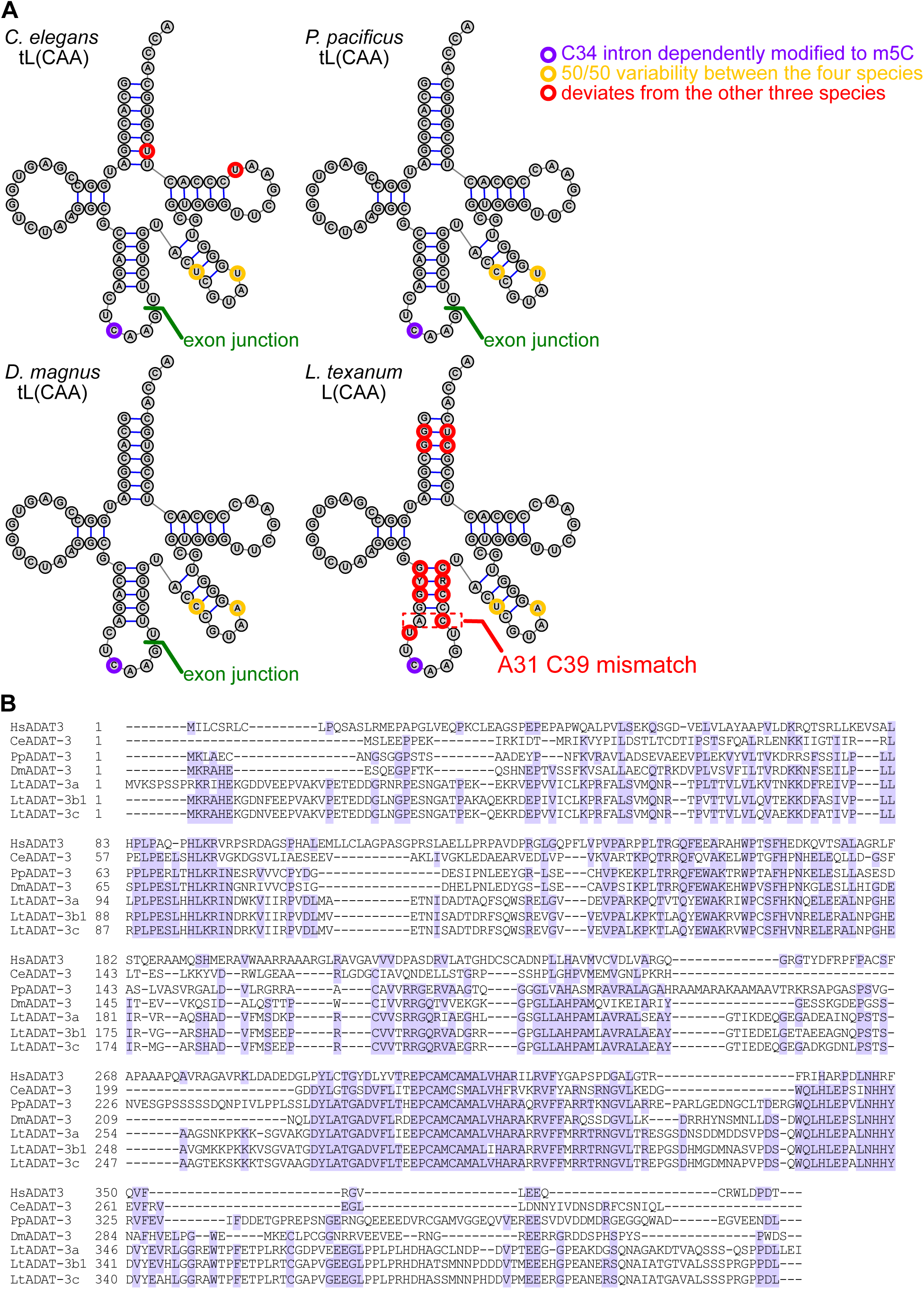
*L. texanum* has an unusual tL(CAA) tRNA and triplicate *adat-3* genes that may relate to the loss of tRNA introns. **A.** Shown are (unmodified) tL(CAA) tRNA sequences in clover-leaf secondary presentation for *C. elegans*, *P. pacificus*, *D. magnus* and *L. texanum*. For the first three, the exon-exon junction resulting from tRNA splicing is indicated in green. C34, which is methylated by NSUN2 in yeast and human, is highlighted in purple. Highlighted in red are residues that differ from the other three sequences. Highlighted in orange are residues that differ in two of the other sequences. All other residues are identical between the four sequences. **B.** Shown is an alignment of ADAT-3 proteins from human, *C. elegans*, *P. pacificus*, *D. magnus*, and *L. texanum*.

The most striking change in the *L. texanum* tL(CAA) genes was a C at position 39, compared to a T in other nematodes (Figure 2A). This mutation is two nucleotides 3’ of the intron position. This T to C change in *L. texanum* disrupts basepairing with A31, which normally forms the bottom basepair of the anticodon stem. Thus, if these bases are not modified, a conserved A-U basepair is replaced by A-C. This is further discussed in the next section.

Another notable change is that the conserved C32 is mutated to T in the *L. texanum* tL(CAA) genes. Position 32 flanks the A-C mismatched base pair (Figure 2). Interestingly, both the nucleotide at position 32 and the intron have been implicated in methylation of C34 in the anticodon. Methylation of C34 by NSUN2 requires the intron in both human (35) and yeast (36). In human tL(CAA), changing C32 to U causes a small decrease in methylation of C34 by NSUN2, while changing the same C to G eliminates it (35). It is possible that the disruption of the 31-39 basepair and/or U32 allows for the intron-independent formation of ^m5^C34 in *L. texanum*, Unfortunately, very little is known about tRNA modification in C. elegans and further experiments would be needed to clarify the modification state of this tRNA in both *C. elegans* and *L. texanum* to determine the relevance of the C32T mutation.

### The L. texanum genome contains a tandem triplication of the *adat-3* RNA deaminase gene

The A31-C39 mismatched basepair can be corrected by RNA deamination. A31 can be deaminated to inosine (I), which can basepair with C39. Alternatively, C39 can be deaminated to U. Both C to U and A to I editing are well known reactions in other aspects of RNA biology, including editing of A34 and A37 in the anticodon of tRNAs (37, 38). However, editing is rarely used to restore base-paring within a tRNA. Precedence for restoring a A-C mismatch in pre-tRNA by de-amination can be found at C4 in plant mitochondria (39). Thus, while restoring the encoded A-C to a Watson-Crick basepair would use well-known chemistry, such a change in the 31-39 basepair would be unprecedented.

The unusual A-C basepair in tL(CAA) of *L. texanum* led us to examine orthologs of RNA deaminases that convert either C to U or A to I. These are known as APOBECs, ADARs and ADATs, which are all related to each other. APOBECs and ADARs are not known to act on tRNA. Furthermore, *C. elegans* does not have an APOBEC, and we were unable to find a homolog in the Diplogasteroidea. ADARs deaminate A to I in long double stranded regions of RNA and the *C. elegans* genome is annotated with two family members, ADR-1 and ADR-2 (40). The *L. texanum* and *D. magnus* genomes each encode single orthologs of both ADR-1 and ADR-2. Thus, *L. texanum* resembles *C. elegans* in its APOBEC and ADAR complement. There are two classes of ADATs (adenosine deaminases acting on tRNA), with ADAT1 modifying A37 next to the anticodon of tRNA Ala in human and yeast (41–43). ADAT1 is absent from *C. elegans* and we were unable to find a homolog in the Diplogasteroidea.

Most interestingly, a heterodimer of ADAT2 and ADAT3 deaminates A34 to I in tRNA (44, 45). A34 to I editing is widespread in eukaryotes and allows a single tRNA to decode both NNU and NNC codons for Ala, Pro, Thr, Val, Ser, Arg, Leu and Ile (Supplemental Figure S1). Within this heterodimer ADAT2 is the catalytic subunit, but requires its partner ADAT3 for activity (44, 45). tRNAs with a GNN anticodon for these amino acids are generally absent from eukaryotes, which is true for *C. elegans*, *P. pacificus* and *L. texanum* (Supplemental Figure S1). ADAT2/3 deaminates the ANN anticodon tRNA to generate INN tRNAs, which can decode the NNC codons. The *C. elegans* and *P. pacificus* and *D. magnus* genomes each encode single orthologs of both subunits (Table 1, Figure 2B). Remarkably, *L. texanum* contains a single gene for *adat-2*, but three *ADAT-3* genes, which we named *adat-3a*, *adat-3b* and *adat-3c* (Table 1; Figure 2B). RNAseq data indicates that *adat-3a* is the most highly expressed, and *adat-3c* has very low expression. Furthermore RNAseq data indicates that *adat-3b* is alternatively spliced to produce two different C-termini, further diversifying the ADAT-3s of *L. texanum*. The *adat-3a*, *adat3b* and *adat3c* genes are located in a 80 kb region of contig 42 (Table 1). The most closely related species, *D. magnus*, has only a single *adat-3* ortholog (Table 1; Figure 2B) This suggests that the *adat-3* gene was triplicated through a segmental duplication in *L. texanum* after it diverged from all other species. Within the ADAT2/3 complex ADAT3 appears to recognize the overall tRNA fold, but does not appear to have much sequence specificity (45). We speculate that the different ADAT-3 subunits of *L. texanum* may confer different substrate specificity onto ADAT-2.

Although ADATs are named for their adenosine deamination, they are also capable of deaminating C to U, and they can act on residues other than the A34 of the anticodon. For example, the trypanosome enzyme can deaminate C32 and A34 on the same tRNA (45). Furthermore, the tRNA binding domain of ADAT-3 is connected by a flexible linker, which may allow one of the *L. texanum* ADAT-3s to present either A31 or C39 to the ADAT-2 active site for deamination (45). Overall, *L. texanum* differs from other nematodes in that it lacks an intron in tL(CAA), its tL(CAA) contains a U39C mutation, and its *adat-3* gene is triplicated. It is tempting to speculate that one of the *adat-3* genes functions to convert A31 or C39 of tL(CAA) to compensate for the U39C mutation.

### The *L. texanum* genome encodes orthologs of intron-dependent tRNA modifying enzymes

Several tRNA modifications are known to be intron-dependent (8, 9), presumably because the modifying enzymes recognize the intron-containing precursor, but cannot recognize the spliced tRNA. This includes methylation of C34 of tL(CAA) by NSUN2 in human and yeast, the conversion of U35 to Ψ35 in tY(GUA) by PUS7 in human, yeast, and plants, and the conversion of U34 and U36 to Ψ in tI(UAU) by PUS1 in animals and yeast (8). The most recently discovered intron-dependent modification is the conversion of A58 to ^m1^A58 in yeast tY(GUA) carried out by a heterotetramer consisting of two catalytic subunits (Trm61) and two noncatalytic subunits (Trm6) (9). It is not yet clear to what extent this intron-dependent activity is conserved for other tRNAs and/or for other species, but it is clear that this enzyme methylates some other tRNAs that don’t have an intron (46).

These intron-dependent tRNA modification enzymes have clear orthologs in *D. magnus* and *L. texanum* (Table 1), and multiple sequence alignments did not reveal any striking differences between the orthologs of species that did or did not have introns in the corresponding tRNA (Supplemental Figure S4 A-E). These enzymes have functions in addition to intron-dependent tRNA modifications. For example, some of them modify snRNAs and mRNAs in addition to tRNAs. Thus, the presence of these enzymes does not predict that any particular tRNA is modified.

### The tRNA ligase complex and *xbp-1* splicing are conserved in *L. texanum*

In addition to TSEN, tRNA splicing requires an RNA ligase (5). In human, tRNA ligase is a complex of five subunits: RTCB, DDX1, FAM98B, CGI99 and ASW (47). Four of these have annotated orthologs in *C. elegans* (Table 1). An ortholog for the fifth subunit, ASW, is not readily apparent because it is small protein with low sequence conservation. This RTCB ligase complex requires activation by another enzyme, archease, which also has a clear ortholog in *C. elegans* (Table 1). In *C. elegans*, RTCB-1 is required for both tRNA and *xbp-1* splicing (48). Using the *C. elegans* tRNA ligase subunits as queries in a BLAST search readily identifies orthologs in both *D. magnus* and *L. texanum* (Table 1). The tRNA ligase subunits and the archease activator were highly similar to the *C. elegans* orthologs (Supplemental Figure S5). Thus, *L. texanum* likely encodes a functional tRNA ligase complex, even in the absence of tRNA introns.

The absence of tRNA introns but the presence of a tRNA ligase complex suggests that the ligase complex may be retained for *xbp-1* mRNA splicing. To determine whether *L. texanum* retained the *xbp-1* splicing, BLAST was used to identify orthologs of the endonuclease IRE-1 that initiates splicing and its *xbp-1* target mRNA in *D. magnus* and *L. texanum*. IRE-1 and XBP-1 orthologs were readily identified (Table 1; Figure S5). The *ire-1* genes are very large, and in the Illumina *D. magnus* and *L. texanum* genome assemblies one half of the gene is on one contig and the other half on another contig. RNAseq reads connected these two halve genes suggesting that the contigs were adjacent to each other in the genome. The HiFi assembly of the *L. texanum* genome confirmed this and revealed a large (>22 kbp) intact *ire-1* gene that encodes for a 1059 amino acid protein.

To identify introns in the *xbp-1* orthologs of *L. texanum* and *D. magnus*, RNA sequencing reads from the Sequence Read Archive (SRR12424051 and SRR12424050) were aligned to the genome (Figure 3A). This revealed that the *xbp-1* genes from *D. magnus* and *L. texanum* contained four and three introns, respectively. Two of the three *L. texanum* introns and three of the four *D. magnus* introns are efficiently spliced and resemble spliceosomal introns, with canonical 5’ and 3’ splice sites (GU and AG). The final intron in both species closely resembles previously described IRE1- and tRNA ligase-dependent introns, with the splice sites occurring in 7-mer CAGCAGN loops of short stem-loop regions (6) (Figure 3A and B). Both the sequence and the structure of this intron are conserved when compared to *C. elegans*, and hundreds of RNA sequencing reads show the intron is indeed spliced out in *L. texanum* and *D. magnus*. Specifically, 153 and 287 reads correspond to the spliced *xbp-1* mRNA in *L. texanum* and *D. magnus,* respectively. These exon-exon junction reads are a minority of the reads under the non-inducing conditions used in these RNAseq experiments, consistent with splicing of this intron being regulated. This analysis clearly indicates conserved splicing of a UPR-type intron in the *xbp-1* mRNA of *L. texanum*. The observation that an *xbp-1*-like intron is conserved in the *L. texanum* ortholog explains why the tRNA ligase complex, archease, and IRE1 are conserved.

**Figure 3:**
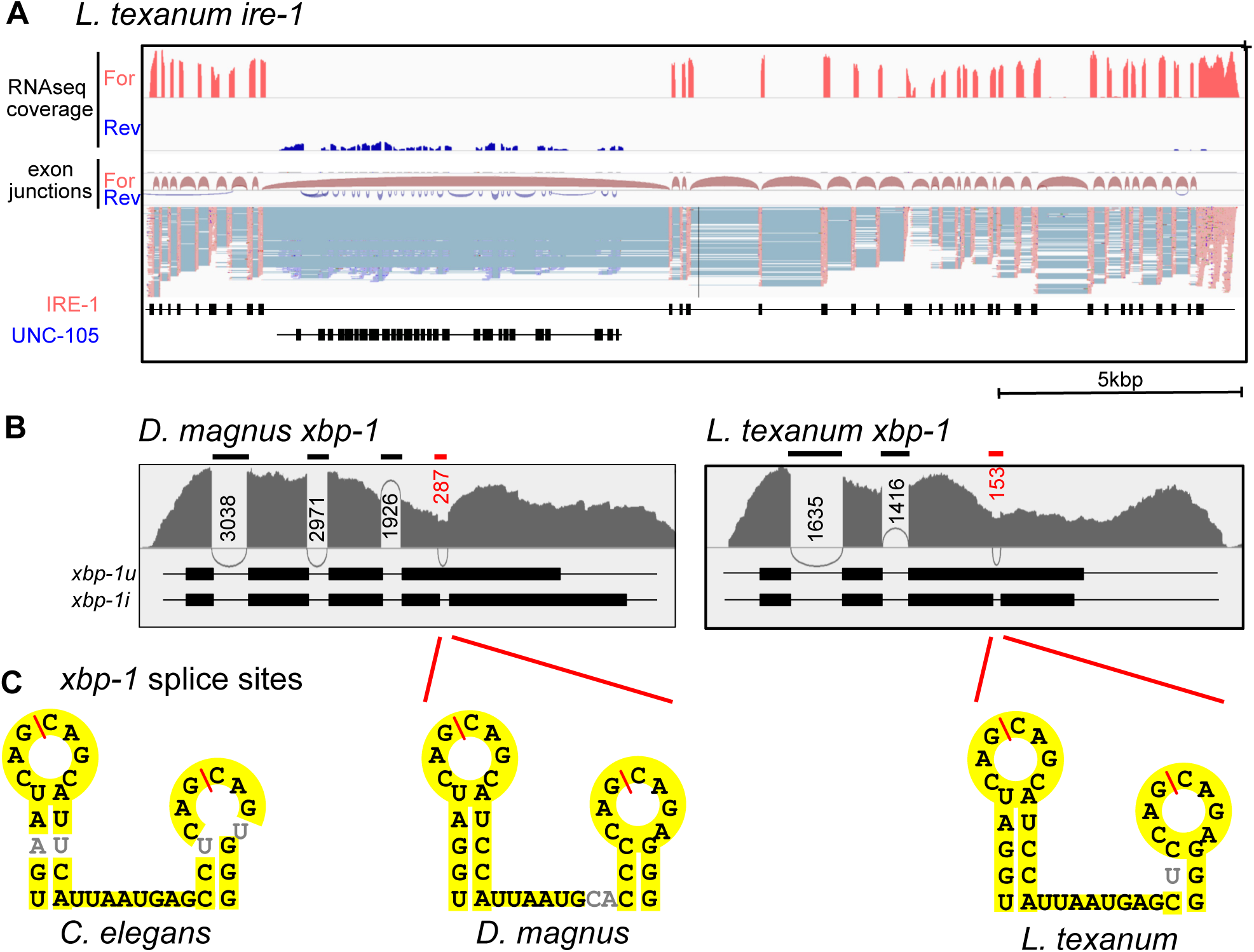
The *xbp-1* gene of *L. texanum* includes an *XBP1*-type intron. **A.** The >22 kbp *ire-1* gene of *L. texanum* identified in a single contig of the HiFi assembly, demonstrating the utility of this improved genome assembly. 35 exons and 34 introns were readily identified using RNAseq data. RNAseq coverage and exon junctions are shown in red for the IRE-1 coding strand and blue for the antisense strand in the top 4 tracks. The fifth track shows alignment of individual reads with the introns in cerulean. The 9^th^ intron is large, and the opposite strand contains the ortholog of *unc-105*, as is the case in *C. elegans*. **B.** Shown are Sashimi plots that depict RNAseq read coverage and exon junction reads along the *xbp-1* orthologs of *D. magnus* (left) and *L. texanum* (right). Spliceosomal introns are marked with a black bar and UPR-type introns are marked with a red bar. Boxes below indicate the coding sequences of the mRNA with the UPR intron retained (*xbp-1u)* and spliced out (*xbp-1s*) **C.** Sequence and predicted secondary structure of the splice sites of the UPR-type introns from *C. elegans, D. magnus,* and *L. texanum* with splice sites indicated with a red slash mark. Nucleotides conserved between the three species are highlighted in yellow.

## CONCLUSIONS

We report here the first known eukaryotic genome that lacks tRNA introns and the tRNA endonuclease required to splice such introns. This discovery complements previous publications that *XBP1*-type and spliceosomal introns are lost in some fungi (13–15, 49). We conclude that none of the three classes of eukaryotic nuclear introns is universally conserved. To the best of our knowledge the L. texanum genome if the only animal genome reported to have lost any of the three intron types.

Interestingly, it has previously been shown that yeast TSEN has an additional but unknown role not connected to tRNA splicing (19, 20). Whether this unknown essential function is conserved in other species is not known. Clearly, this function is not conserved in *L. texanum*. In contrast, orthologs of CLP1 and the tRNA ligase complex are conserved in *L. texanum*, and future studies on *L. texanum* may be effective in more precisely defining the tRNA intron-independent functions of these proteins. Similarly, the recently proposed capacity of yeast tRNA introns to base pair to mRNAs and regulate their expression (7) also is clearly not conserved in *L. texanum*.

The most widespread introns occur in tY(GUA), tI(UAU), and tL(CAA), and in each case these introns are required for modifications in the anticodon by PUS7, PUS1, and NSUN2, respectively, and outside the anticodon by Trm61/Trm6 (8, 9). All four of these intron-dependent tRNA modification enzymes have clear orthologs in *L. texanum* (Table 1 and Supplemental Figure S4). Each of these enzymes also has additional functions, including modifying other tRNAs in an intron-independent manner, and modifying other ncRNAs and mRNAs. It will be interesting to characterize the modification state of these tRNAs in *L. texanum* and the specificity of its ADAT2/3, PUS-1, PUS-7, NSUN-2, and TRMT-61/TRMT-6 orthologs. Unfortunately, very little is known about the modification state of any nematode tRNA, which would also be required to interpret any findings of tRNA modification in *L. texanum*.

These and other future studies would require development of experimental tools for *L. texanum*. Experiments in *C. elegans* are facilitated by robust RNAi in response to feeding dsRNA (50). This experimental approach requires the *sid-1* and *sid-2* genes (51, 52). The *L. texanum* genome does not include *sid-1* or *sid-2* genes, so this experimental approach is unlikely to be available in *L. texanum*. However, *C. briggsae* becomes amenable to RNAi feeding upon transgenic expression of *C. elegans sid-2*, which enables the uptake of extracellular RNA (53), and we anticipate that this approach might also be successful in *L. texanum*. Alternatively, CRISPR/Cas9 genome editing has been established in *P. pacificus* (54, 55) and should be readily adaptable to *L. texanu*m.

## METHODS

### *L. texanum* strain and maintenance

*L. texanum* strain TMG5 was kindly provided by Dr. Erik Ragsdale (Indiana University). Worms were maintained according to standard *C. elegans* cultivation protocols on nematode growth medium (NGM) plates seeded with *E. coli* strain OP50 as a bacterial food source.

### Preparation of *L. texanum* and genome sequencing

Five NGM/OP50 plates were each inoculated with approximately ten embryos and incubated for six days at 20°C until confluent populations with mixed developmental stages were obtained. Nematodes were washed from these plates with M9 buffer, and approximately 1–3 drops of worm suspension were transferred to fresh plates using a Pasteur pipette. These plates were cultured for an additional 3 days at 20°C to expand the population. Nematodes were harvested by washing 43 plates with cold 0.1 M sodium chloride solution into a 50 ml conical tube. Nematodes were pelleted by centrifugation at 1,000 rpm (r=168mm; 190g) for 1 min. The supernatant was discarded, and the pellet was resuspended in 20 ml of cold 0.1 M sodium chloride solution; this washing step was repeated twice more. After final washing, nematodes occupied approximately 1 ml wet volume and were adjusted to a total volume of 20 ml using cold 0.1 M sodium chloride. 20 ml of cold 60% sucrose solution was carefully layered underneath the worm suspension, followed by an additional overlay of 8 ml cold 0.1 M sodium chloride solution. The gradient was centrifuged at 2,000 rpm (r=168mm; 760g) for 3 min to separate nematodes from debris and bacteria, to minimize the presence of *E. coli* DNA. The nematode layer was collected, diluted into 15 ml of cold sterile water, and pelleted again by centrifugation at 1,000 rpm for 1 min. This wash was repeated twice more. Following the final wash, nematodes were concentrated into ∼500 µl of cold sterile water and transferred into microcentrifuge tubes. Approximately 300 µl total wet worm pellet volume was flash-frozen on dry ice and stored at −80°C until shipment to CD genomics (Shirley, NY 11967) for high molecular weight DNA extraction and PacBio HiFi sequencing. DNA sequencing reads were deposited in the sequence read archive under accession number PRJNA1267722. 624,139 reads totaling 4.57Gbp were obtained and used in genome assembly with hifiasm (31). Contigs corresponding to the mitochondrial genome were removed by comparison to the *P. pacificus* mitochondrial DNA (Genbank NC_015245.1) with MitoHiFi (56). Contaminating contigs (from OP50) were removed using Kraken2 (57) and duplicate contigs were removed with funannotate assembly clean (https://zenodo.org/records/4054262). The genome assembly was deposited in Genbank under accession number PRJNA1267722.

### tRNA annotation and alignment

Genome sequences for *L. texanum* (GCA_014805385.1), *D. magnus* (GCA_014805445.1), *M. japonica* (GCA_900490955.1)*, P. pacificus* (GCA_000180635.4)*, P. giblindavisi* (GCA_943737045.1)*, A. sudhausi* (GCA_945643635.2), *and K. luziae* (GCA_014805365.1) were downloaded from NCBI (https://www.ncbi.nlm.nih.gov/) and analyzed by tRNAscan-SE 2.0 in the eukaryotic mode. tRNAscan-SE 2.0 was also used to predict tRNA genes in the HiFi *L. texanum* assembly. The analysis in the main text and Figure 1 is on the hifiasm assembly, but the Illumina based GCA_014805385.1 assembly supported the same conclusions. The tRNAscan-SE program gives each prediction a score with scores below 55 bits indicating a likely pseudogene. The only candidate tRNA intron predicted for *L. texanum* was in a possible tL(CAG), but this is very likely a pseudogene: The algorithm predicted 25 possible genes for tL(CAG), of which 20 were identical and scored very high. The gene with a possible intron received the lowest score (41.25), due to two mismatches that would interfere with formation of the anticodon stem and a mismatch in the anticodon loop. This prediction was therefore discarded. Predicted tRNA sequences were aligned with Clustal and depicted as sequence logos in the supplemental Figure S3.

### Identification of tRNA splicing endonuclease, tRNA modification, tRNA ligase genes, and *ire-1* genes

The protein sequences for human proteins listed in Table 1 were retrieved from Genbank and used to search *C. elegans* annotated proteins by BLASTP. No annotated homologs of TSEN15 or ASW could be identified. Searching the *C. elegans* genome by TBLASTN also failed to identify these genes. A gene for TSEN15 is annotated for the fruit fly genome, but using that in a PSIBLAST search also failed to identify a *C. elegans* homolog. ASW and TSEN15 are both small proteins that lack a catalytic site or other highly conserved sequences and thus may be too far diverged to be identifiable based on sequence similarity. Alternatively, it is possible that in nematodes these proteins are not required to form a functional enzyme complex.

To identify *P. pacificus*, *L. texanum* and *D. magnus* orthologs, the *C. elegans* proteins were used as queries in TBLASTN searches against the genome in Genbank and BLASTP searches against annotated proteins in https://parasite.wormbase.org/ and http://www.pristionchus.org/blast/. Both approaches identified the same genes/genomic regions. After assembly of the HiFi genome, it was also similarly searched using TBLASTN, and the same genes were found. Finally, a preliminary annotation of coding sequences derived from the PacBio assembly was searched with BLASTP, which also found the same genes.

Analyzing RNAseq data, described below, revealed that some of the exons and introns were misannotated in https://parasite.wormbase.org/ or http://www.pristionchus.org/blast/. In the Illumina assemblies, some gene annotations at the end of a contig were missing a part of the protein. This was evident from RNAseq reads that partially mapped to these contigs. The other (Non-mapping) part of these RNAseq reads were used in a BLASTN search to identify the contig that contained the remainder of the gene. These contigs similarly had some RNAseq reads that partially mapped to them and partially to the original contig. Furthermore, the breakpoint of mapping reads resembled a spliceosomal intron. This strongly suggest that the two contigs are next to each other in the genome and that there is a unsequenced gap in the intron. All of the genes of interest were contained in a single contig in the HiFi assembly. The protein sequences were corrected based on the exons and introns evident from RNAseq alignments before conducting the multiple protein sequence alignments by MUSCLE.

### Identification of UPR type introns in XBP-1 orthologs

The XBP-1 orthologs of *L. texanum* and *D. magnus* were identified by BLAST as described above. To identify potential UPR-type introns, RNAseq reads were downloaded from Genbank (SRR12424050 for *L. texanum* and SRR12424051 for *D. magnus*) and aligned to the genome using HISAT2 and RNA STAR (58, 59). Introns were defined by RNAseq reads that spanned the exon-exon junction. Both aligners gave essentially identical results. The sashimi plots of Figure 3B were generated using IGV.

## ACKNOWLEDGEMENTS

We thank Erik Ragsdale (Indiana University) for sharing the *L. texanum* strain, Anita Hopper (Ohio State University) and the van Hoof lab for stimulating discussions, and Catherine Stuart for editing the manuscript. This work was funded by NIH grants R35GM141710 to AvH and R35GM140933 and R01HD101269 to S.A.

## SUPPLEMENTAL RESULTS AND DSICUSSION

### The tRNA variety of *L. texanum* and *P. pacificus* is identical

The loss of introns from the *L. texanum* tRNA genes does not seem to correlate with other changes in the tRNA gene pool (Figure S1). We detected 48 different tRNA types in the HiFi *L. texanum* assembly and in the *P. pacificus* genome (https://gtrnadb.ucsc.edu/genomes/eukaryota/Ppaci4/) (1, 2). The same 48 tRNAs were present in the Illumina *L. texanum* assembly, although there were slight variations in the number of genes for some tRNA types.

The initial tRNAscan-SE analysis detected 46 tRNA types. The 47^th^ type is an unusual Leu tRNA. One striking feature of the *C. elegans* tRNA pool is that it has two different tRNAs with UAU anticodons: it has a typical Ile tRNA with a UAU anticodon, but it also has an unusual Leu tRNA with a UAU anticodon (1, 2). This tL(UAU) tRNA (highlighted with a red outline in Supplemental Figure S1) is classified as tI(UAU) by tRNAscan-SE, but the two tRNAs are readily distinguished by their sequence as well as the longer variable arm and overall length of the Leu tRNA. Both *P. pacificus* and *L. texanum* (Supplemental Figure S1) also had these two different tRNAs.

The 48^th^ type comes from separating two different Met tRNAs. The tRNAscan-SE output groups initiator Met tRNA and elongator Met tRNAs as one isodecoder even though they have distinct functions. We curated the Met tRNAs based on sequence similarity to *C. elegans* and *P. pacificus* tRNAs.

Overall, this analysis revealed a typical repertoire of tRNA genes for *L. texanum* (Supplemental Figure S1). The only difference in tRNA types that we detected is that *L. texanum* and *P. pacificus* have three different glycine tRNAs, while *C. elegans* only has two (Supplemental Figure S1). This indicates that in *C. elegans*, tG(UCC) decodes both GGA and GGG codons, while in *P. pacificus* and *L. texanum*, tG(CCC) may contribute to the decoding of the GGG codons. The third Glycine tRNA is highlighted with a green outline in Supplemental Figure S1. We do not think that this change is related to the loss of introns.

The total number of tRNA genes per eukaryotic genome is highly variable, and consistent with this, the HiFi *L. texanum* genome contains approximately three times as many tRNA genes as *C. elegans* and about 50% more than *P. pacificus* (Supplemental Figure S1). Strikingly, *L. texanum* has a drastically increased number of tC(GCA) genes (highlighted with an orange outline in Supplemental figure S1). The difference is explained by *L. texanum* contig 52, which contains 413 copies of tC(GCA) in a 84,056 bp stretch. These 413 tandem repeats of a single tRNA almost completely account for the 556 increase in *L. texanum* tRNA gene number compared to *P. pacificus*. The sequences of the tC(GCA) genes in the repeat where the same as the genes dispersed outside of the repeat. One notable difference between the previous Illumina based *L. texanum* genome assembly and our HiFi assembly is that the tC(GCA) repeat was missing from the Illumina assembly because of the difficulty in assembling such highly repeated regions from small reads. However, the *P. pacificus* assembly is also a high quality PacBio HiFi assembly and thus the difference in tC(GCA) gene number is likely a biological difference and not an assembly quality difference. We do not think that this change in tC(GCA) gene number is related to the loss of introns. Therefore, we conclude that the loss of tRNA introns has no obvious connection to the tRNA pool.

## SUPPLEMENTAL FIGURES LEGENDS

**Supplemental Figure S1:**
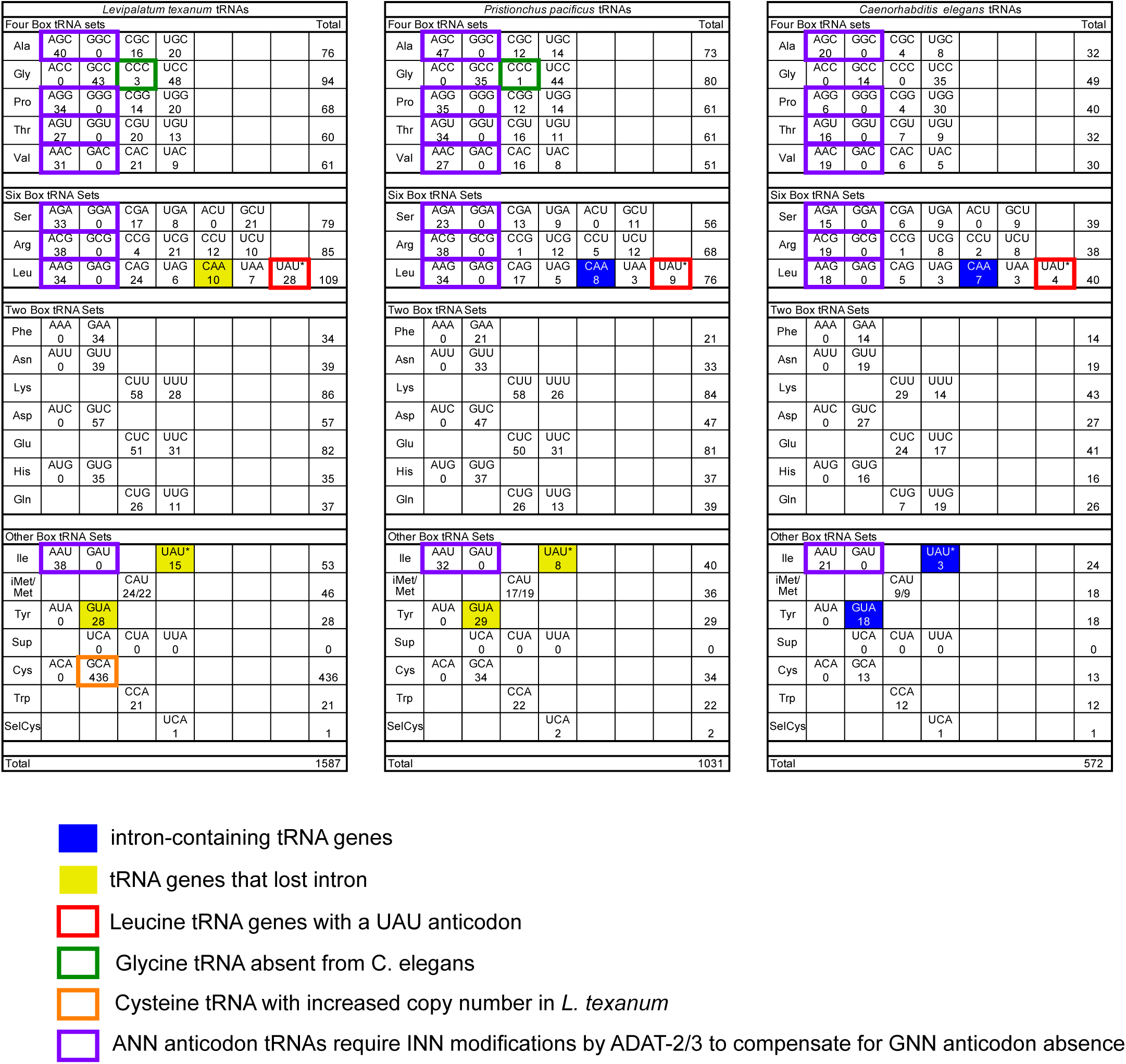
tRNA genes in the *L. texanum* genome compared to those in *P. pacificus* and *C. elegans*. Indicated is the number of genes for each of 48 different tRNAs detected in the HiFi assembly of the *L. texanum* genome (left) *P. pacificus* (middle) and *C. elegans* (right). tRNA genes that contain introns are highlighted in blue and those that have lost introns are in yellow (as in Figure 1). Red outline highlights an unusual leucine tRNA with a UAU anticodon that was previously described in C. elegans and P. pacificus. Green outline highlights tG(CCC) genes present in *L. texanum* and *D. magnus* but not in *C. elegans*. Orange outline highlights the dramatically increased copy number for tC(GCA) in *L. texanum*. Purple outline highlights highlights ANN/GNN anticodon pairs were the GNN anticodon is commonly missing from eukaryotic tRNAs and the ANN anticodon tRNA is modified to INN by ADAT-2/ADTA-3. L. texanum has the typical 8 INN anticodon tRNAs, which does not explain the triplication of *adat-3*.

**Supplemental Figure S2:**
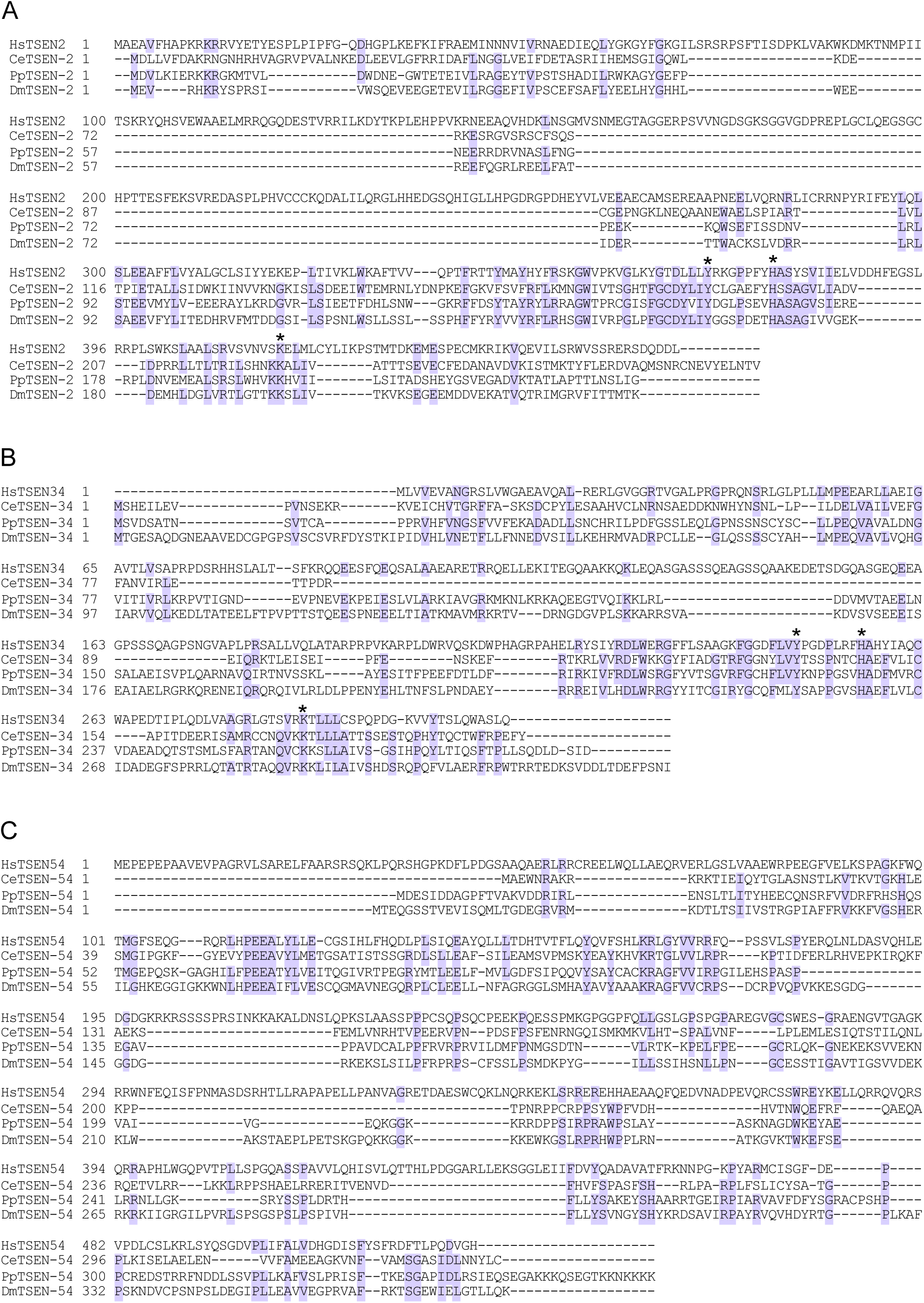
TSEN-2, TSEN-34 and TSEN-54 are conserved in *P. pacificus* and *D. magnus*. Sequences for TSEN-2 **(A)**, TSEN-34 **(B)** and TSEN-54 **(C)** were identified by BLAST and aligned with MUSCLE. Purple shading indicates identity to the consensus sequence. The TSEN subunits show low sequence similarity across eukaryotes but the catalytic residues (indicated with asterisks) are conserved.

**Supplemental Figure S3:**
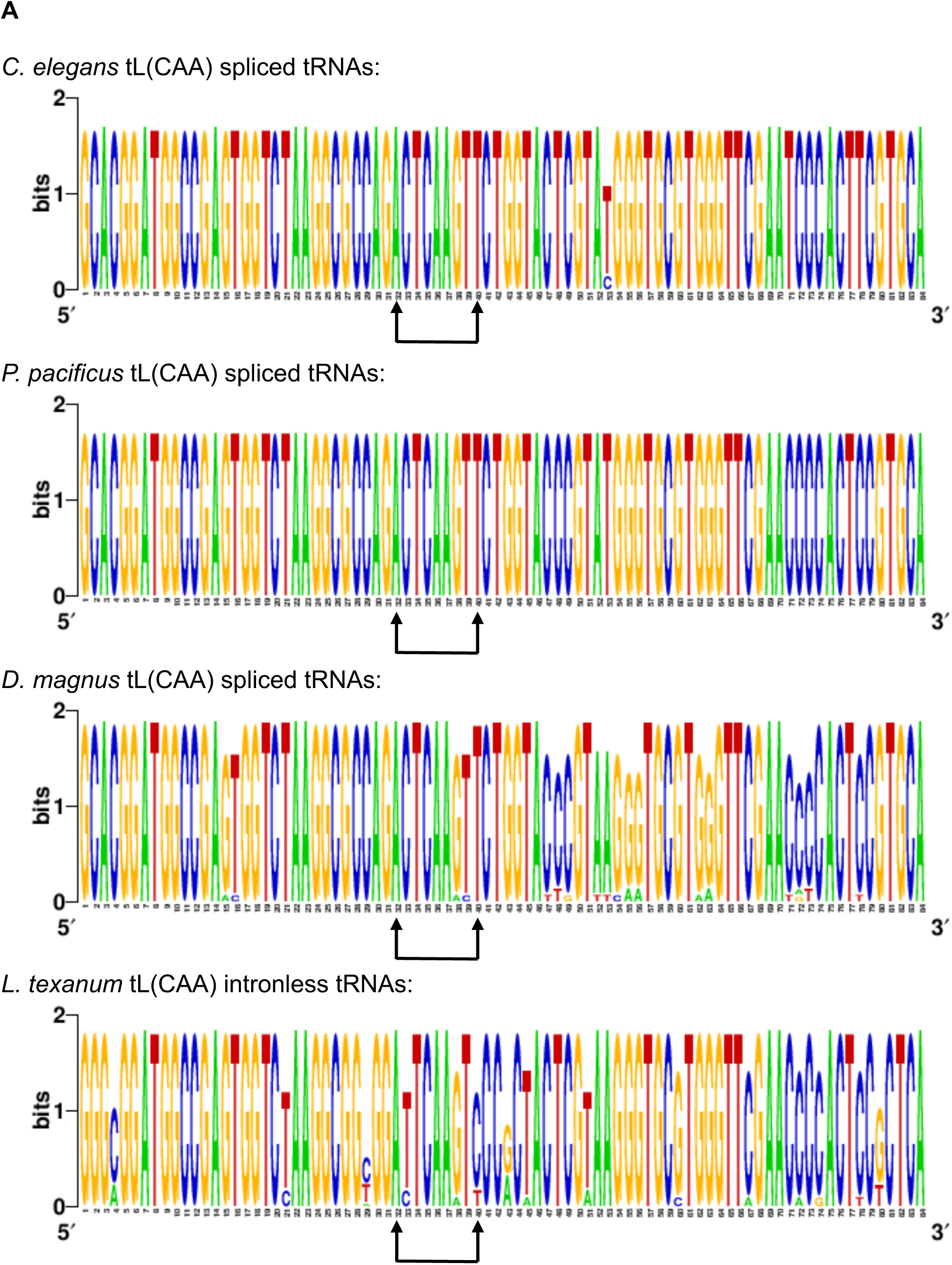
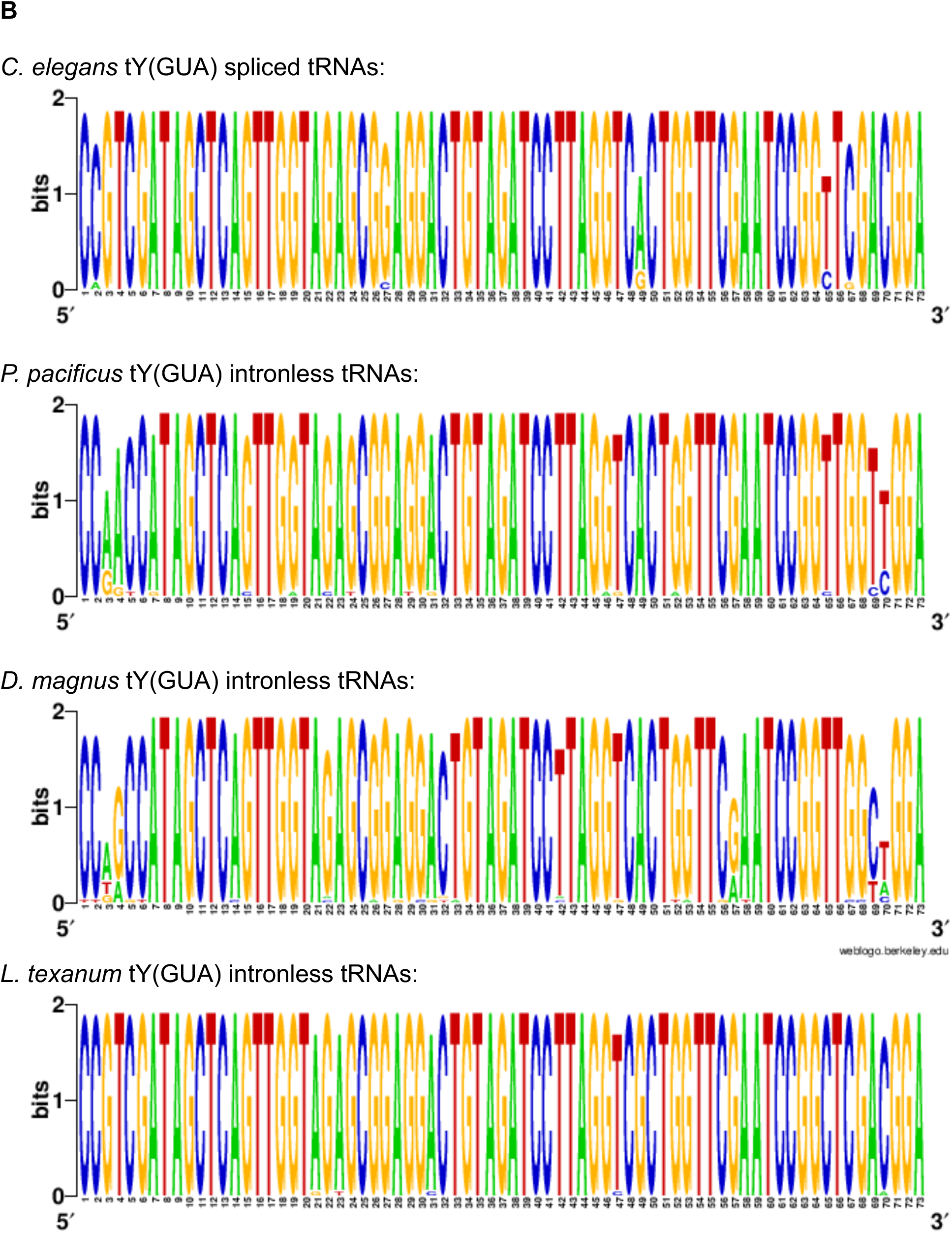
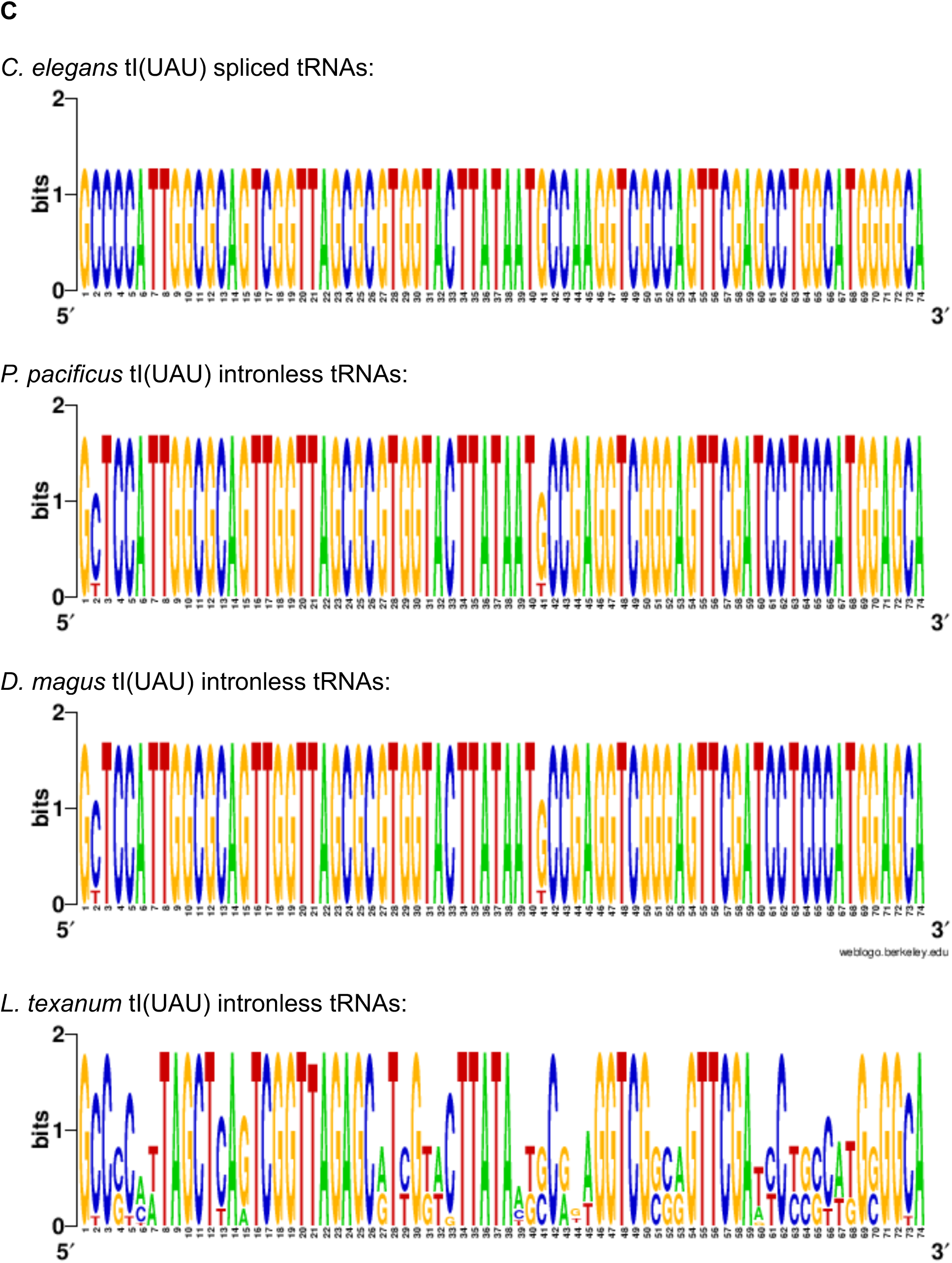
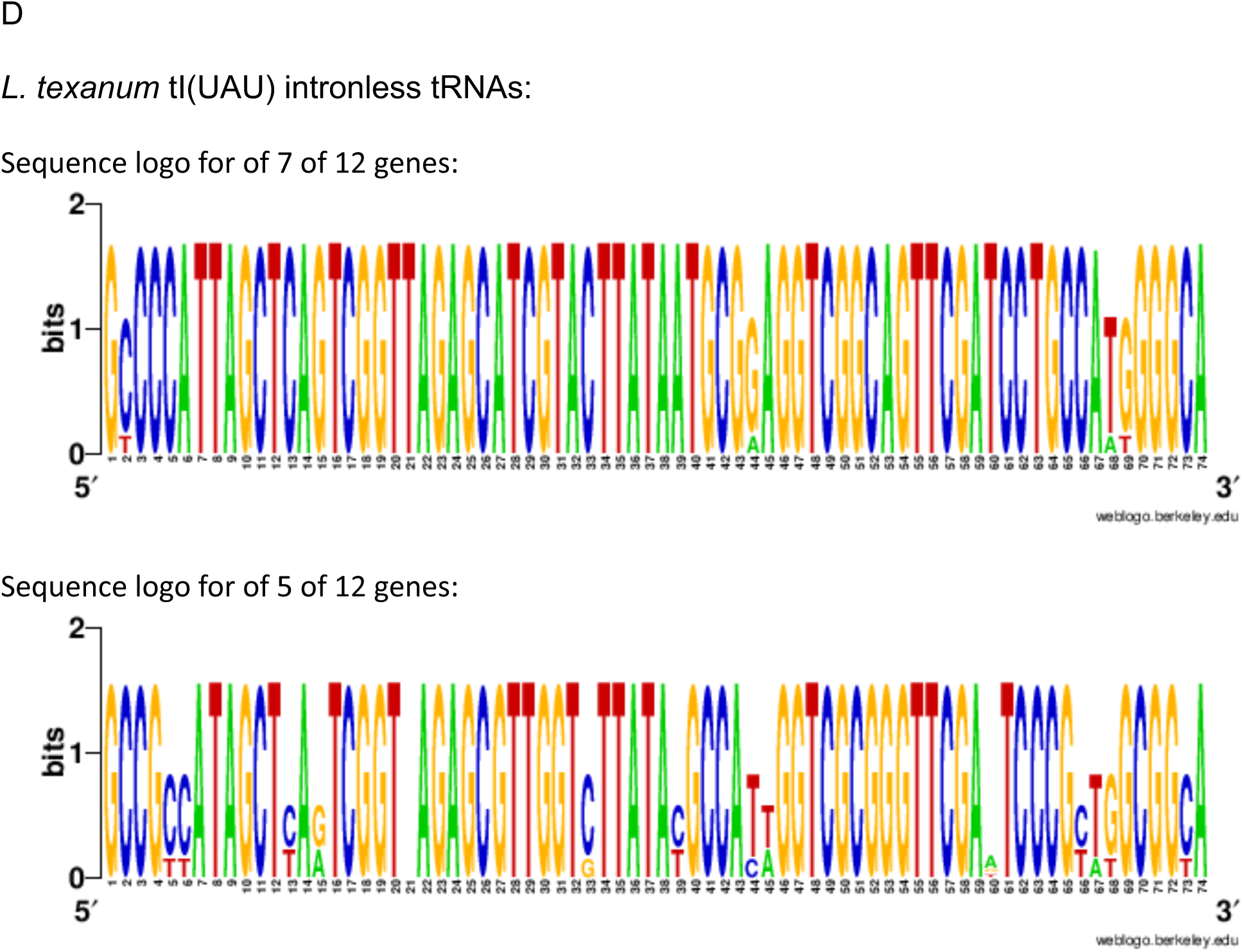
Sequence of tL(CAA), tY(GUA) and tI(UAU) tRNAs in *C. elegans*, *P. pacificus, D. magnus and L. texanum.* Aligned tRNA gene sequences identified by tRNAscanSE and with the introns removed are depicted as sequence logos. (**A)** The tL(CAA) sequence is highly conserved between *C. elegans*, *P. pacificus*, and *D. magnus* but more diverged in *L. texanum* as also depicted in Figure 3. The base pair between positions 31 and 39 is indicated with a double headed arrow. (**B)** The tY(GUA) sequence is highly conserved between the four species. **(C)** The tI(UAU) sequence is highly conserved between *C. elegans*, *P. pacificus*, and *D. magnus* but more variable in *L. texanum*. The divergence in *L. texanum* appears unconnected to the loss of the intron because intron loss is shared with *P. pacificus* and *D. magnus*. **(D)** The variability in tI(UAU) sequence in *L. texanum r*eflects the presence of two gene families. It is unknown whether both families are functional.

**Supplemental Figure S4:**
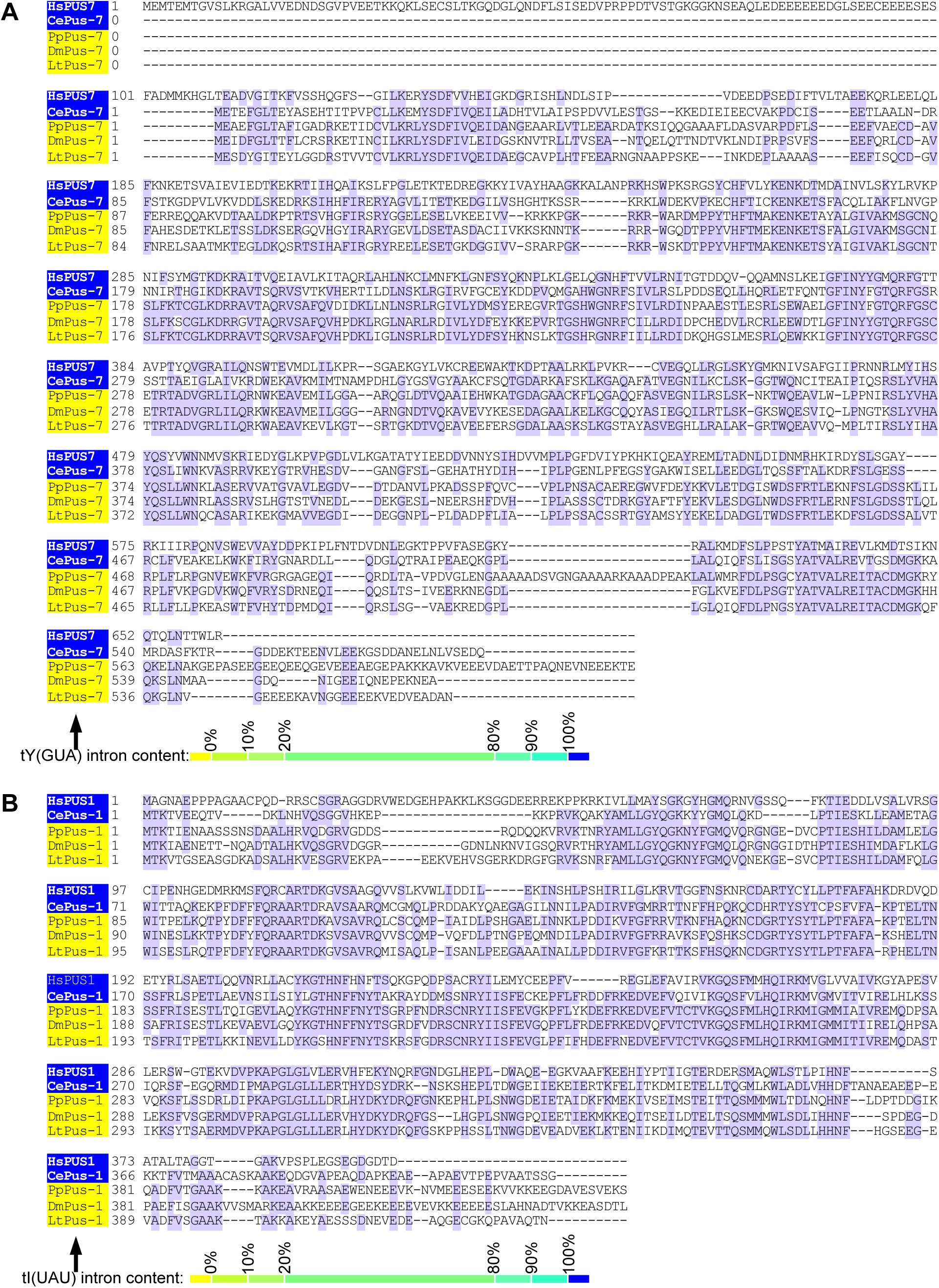
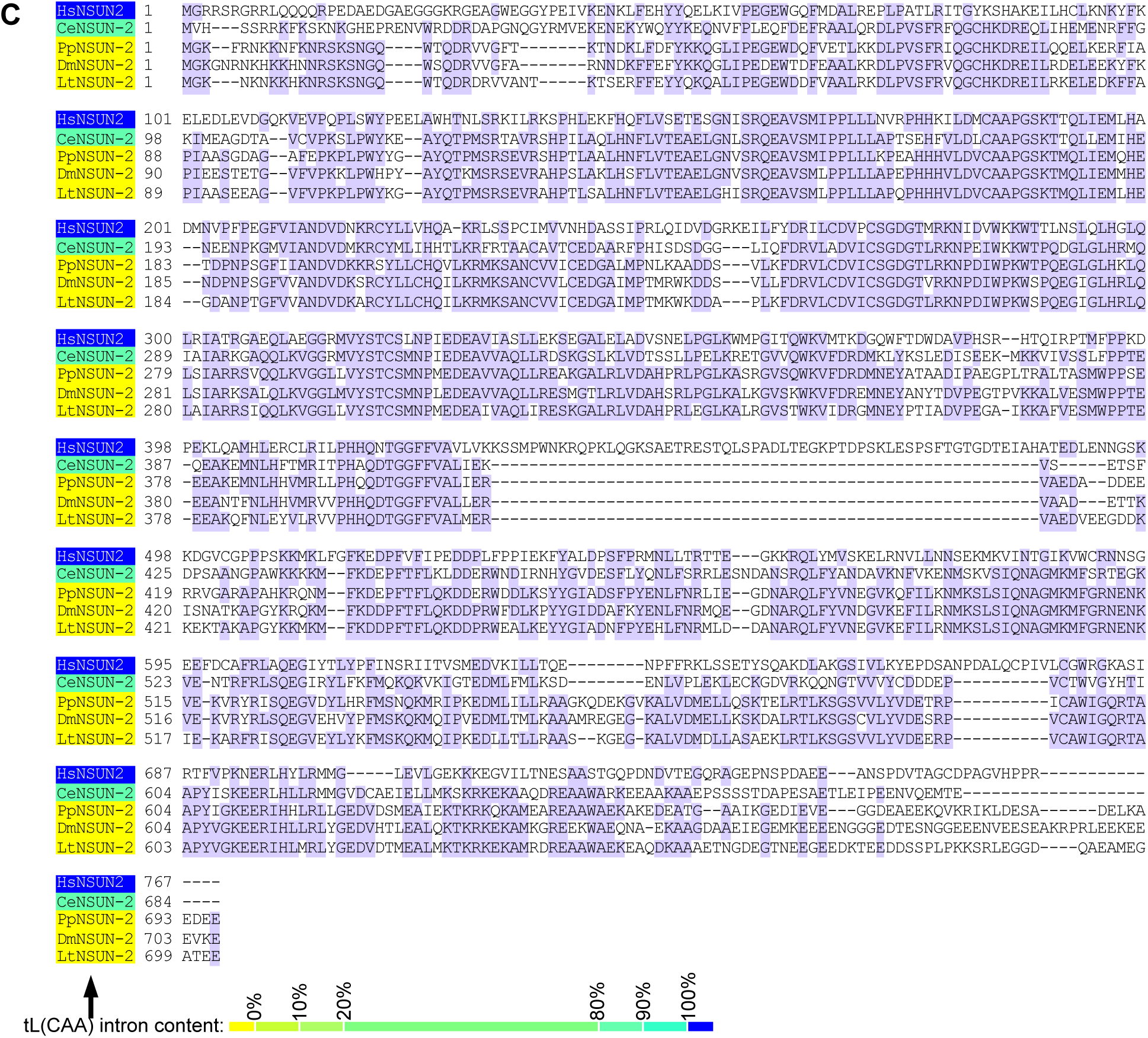
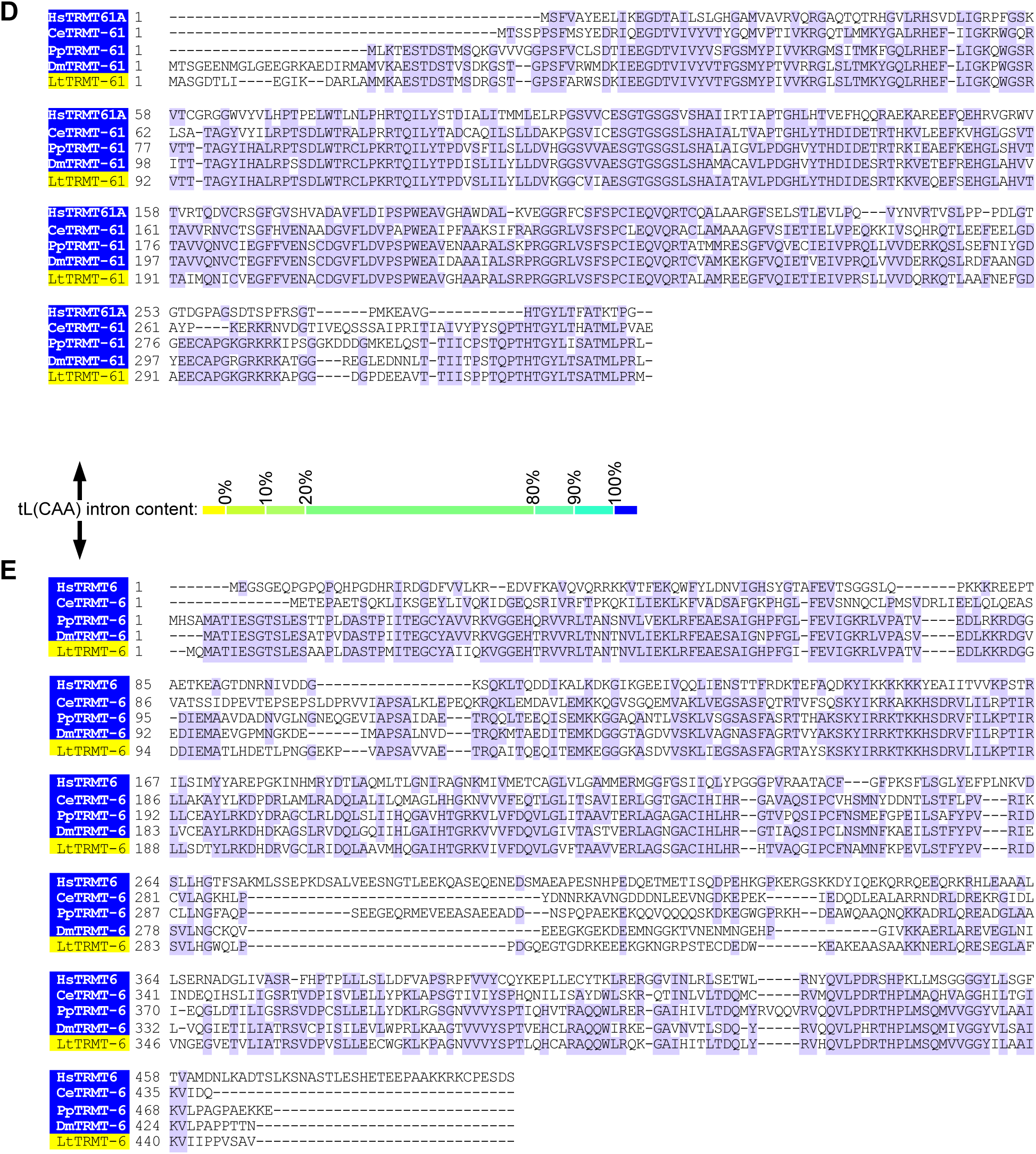
Sequence conservation of intron-dependent tRNA modification enzymes PUS7 (A), PUS1 (B), NSUN2 (C), TRMT61 (D), and TRMT6(E). Sequences were identified by BLAST and aligned with MUSCLE. Purple shading indicates identity to the consensus sequence. Blue and yellow shading reflects intron presence in the indicated tRNA substrates.

**Supplemental Figure S5:**
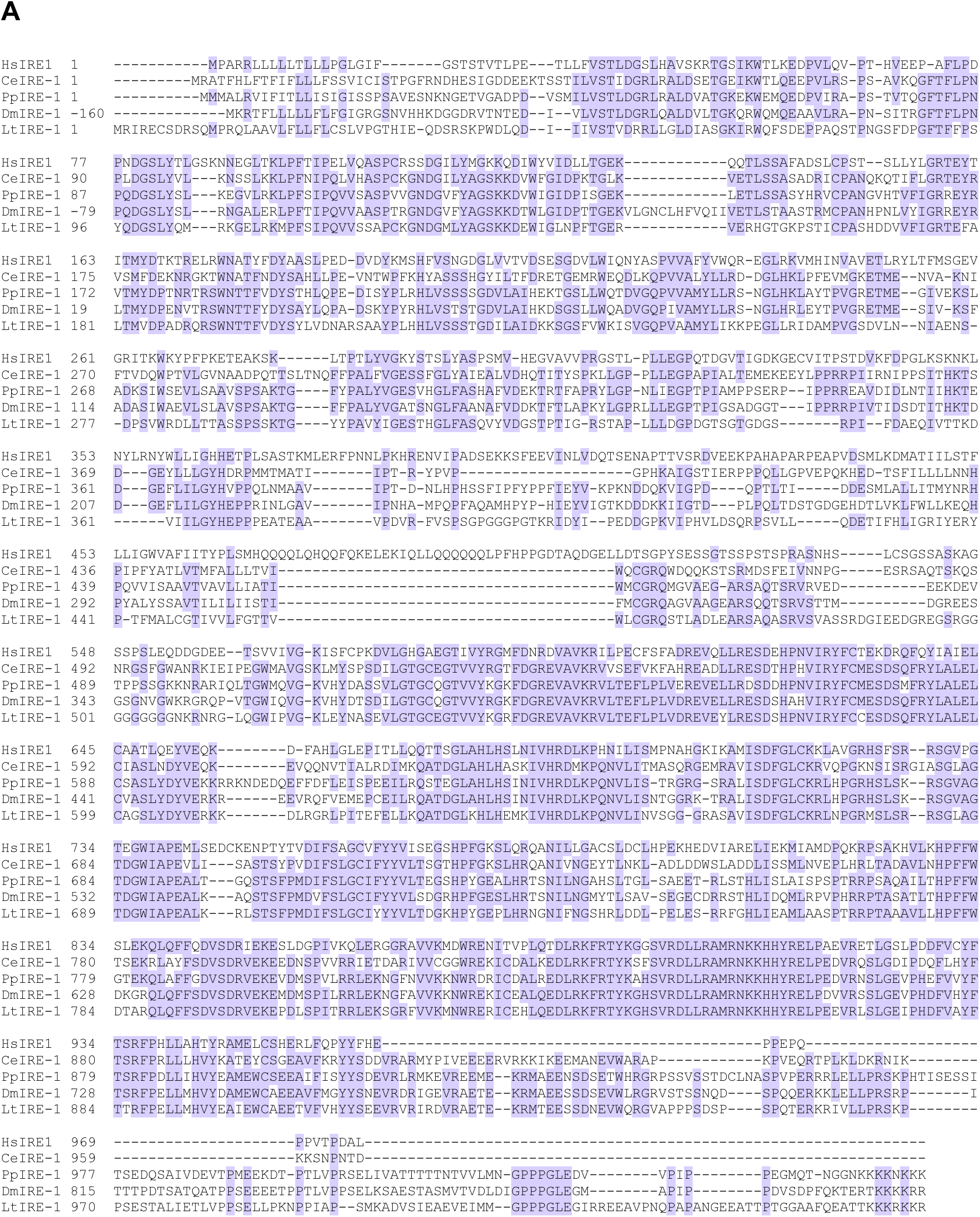
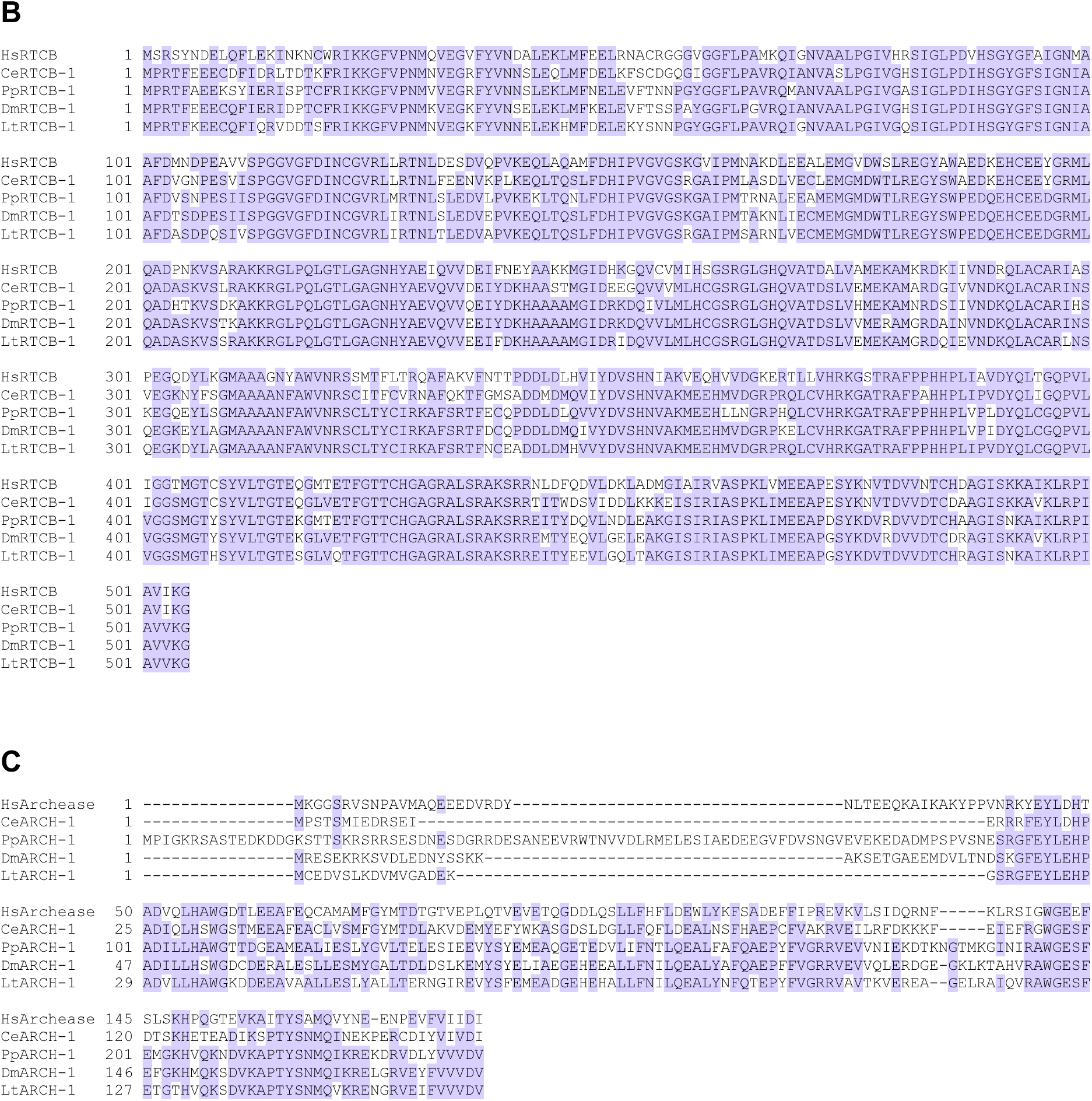
Sequence conservation of *xbp-1* splicing enzymes IRE-1, RCTB-1, and Archease. Sequences for IRE-1 **(A)**, RTCB-1 **(B)** and Archease **(C)** were identified by BLAST and aligned with MUSCLE. Purple shading indicates identity to the consensus sequence.

## Notes

### Competing Interest Statement

The authors have declared no competing interest.

